# MODE-TASK: Large-scale protein motion tools

**DOI:** 10.1101/217505

**Authors:** Caroline Ross, Bilal Nizami, Michael Glenister, Olivier Sheik Amamuddy, Ali Rana Atilgan, Canan Atilgan, Özlem Tastan Bishop

**Author notes:** Equal contribution (Caroline Ross, Bilal Nizami).

## Abstract

**Summary:** MODE-TASK, a novel software suite, comprises Principle Component Analysis, Multidimensional Scaling, and t-Distributed Stochastic Neighbor Embedding techniques using molecular dynamics trajectories. MODE-TASK also includes a Normal Mode Analysis tool based on Anisotropic Network Model so as to provide a variety of ways to analyse and compare large-scale motions of protein complexes for which long MD simulations are prohibitive.

**Availability and Implementation:** MODE-TASK has been open-sourced, and is available for download from https://github.com/RUBi-ZA/MODE-TASK, implemented in Python and C++.

**Supplementary information:** Documentation available at http://mode-task.readthedocs.io.

## 1 Introduction

Conventional analysis of molecular dynamics (MD) trajectories may not identify global motions of macromolecules. Normal Mode Analysis (NMA) and Principle Component Analysis (PCA) are two popular methods to quantify large-scale motions, and find the “essential motions”; and have been applied to problems such as drug resistant mutations (Nizami *et al.*, 2016) and viral capsid expansion (Hsieh *et al.*, 2016).

MODE-TASK is an array of tools to analyse and compare protein dynamics obtained from MD simulations and/or coarse grained elastic network models. Users may perform standard PCA, kernel and incremental PCA (IPCA). Data reduction techniques (Multidimensional Scaling (MDS) and t-Distributed Stochastics Neighbor Embedding (t-SNE)) are implemented. Users may analyse normal modes by constructing elastic network models (ENMs) of a protein complex. A novel coarse graining approach extends its application to large biological assemblies.

## 2 Methods

### 2.1 Implementation

MODE-TASK was developed using C++ and Python programming languages on Linux/Unix-based systems. Non-standard Python libraries, including NumPy, SciPy, Matplotlib (Hunter, 2007), MDTraj (McGibbon *et al.*, 2015) and Scikit-learn (Pedregosa *et al*., 2011), were used. The C++ ALGLIB (www.alglib.net) was also utilized. MODE-TASK supports a wide variety of MD trajectory and topology formats including binpos (AMBER), LH5 (MSMBuilder2), PDB, XML (OpenMM, HOOMD-Blue), .arc (TINKER), .dcd (NAMD), .dtr (DESMOND), hdf5, NetCDF (AMBER), .trr (Gromacs), .xtc (Gromacs), .xyz (VMD), .mdcrd (AMBER) and LAMMPS.

### 2.2 Algorithms

PCA requires the covariance/correlation matrix of atomic positions, obtained from an MD simulation (David and Jacobs, 2014). Modes (eigenvectors) are found by diagonalising the covariance/correlation matrix. Besides standard PCA, MODE-TASK implements memory efficient version (IPCA) through scikit-learn Python library and the original algorithm (Pedregosa et al., 2011; Ross et al., 2008), and different choices of Kernel PCA, a nonlinear generalization of PCA, on an MD trajectory.

The anisotropic network model (ANM) (Atilgan *et al.*, 2001) gives an elastic network model of the protein using all pairs of nodes separated by a defined cutoff. The algorithm builds the Hessian matrix on the atomic coordinates of the C_α_ or C_β_ atom of each residue of a given PDB. It uses the ALGLIB library to calculate the pseudoinverse of the Hessian, leading to the eigenvalues and eigenvectors of the normal modes. A novel coarse graining algorithm is used. An initial C_α_/C_β_ atom is set from a starting residue defined by the user. The distance between this atom and every other C_α_/C_β_ in a single asymmetric unit of the assembly is found. For a specified level of coarse graining, the algorithm selects the n^th^ closest C_α_/C_β_ to the starting point and defines a minimum distance as the distance between the initial and the selected nth closest atom. The algorithm then expands outwards and iteratively steps through the (n+1)^th^ closest atom.

Metric and nonmetric types of MDS for the MD trajectory are implemented using the scikit-learn library. t-SNE is another dimensionality reduction for data of high dimensions (van der Maaten and Hinton, 2008). The dissimilarity measures used are the Euclidean distance between internal coordinates, and pairwise RMSD between the MD frames.

## 3 Performance

Tests were conducted on a PC running Ubuntu 16.04.2 LTS on an Intel Core i7-4790 CPU with a clock speed of 3.60GHz with 32GB of physical memory.

| NMA scripts | Main parameters | Time |
| --- | --- | --- |
| coarseGrain.py | Coarse grain level 4; starting at atom 3 | < 1 sec |
| ANM | Cut off 50 Å; 2460 nodes | 116mins |
| getEigenvector.py |  | < 1 sec |
| conformationMode.py |  | 26 secs |
| Essential dynamics scripts | Main parameters | Time |
| pca.py | SVD solver; 10,000 frames | 26 secs |
| pca.py | RBF kernel; 10,000 frames | 40 mins |
| internal_pca.py | Phi angles; 1,000 frames | 10 secs |
| mds.py | RMSD; 10,000 frames | 88 mins |
| tsne.py | 10,000 frames | 68 mins |

## 4 Applications

### NMA

Coarse graining (level 4, start at residue 3) was performed along the C_β_ atoms (C_α_ for glycine) of the CAV-16 viral capsid (PDB: 5C4W; diameter ≈ 300 Å). The *ANM.cpp* then obtained the eigenvalues and eigenvectors of the respective modes. The cutoff distance was increased to 50 Å (default 15 Å) because of the capsid’s internal cavity. During host cell infection, the capsid expands for RNA-release (PDB: 4JGY). We used *conformationMode.py, getEigenVector.cpp* and *visualiseVector.py* to identify normal modes associated with the structural change. The mode with the largest overlap (0.86) to the conformational change is presented as a radial expansion (Figure 1A).

**Figure 1.**
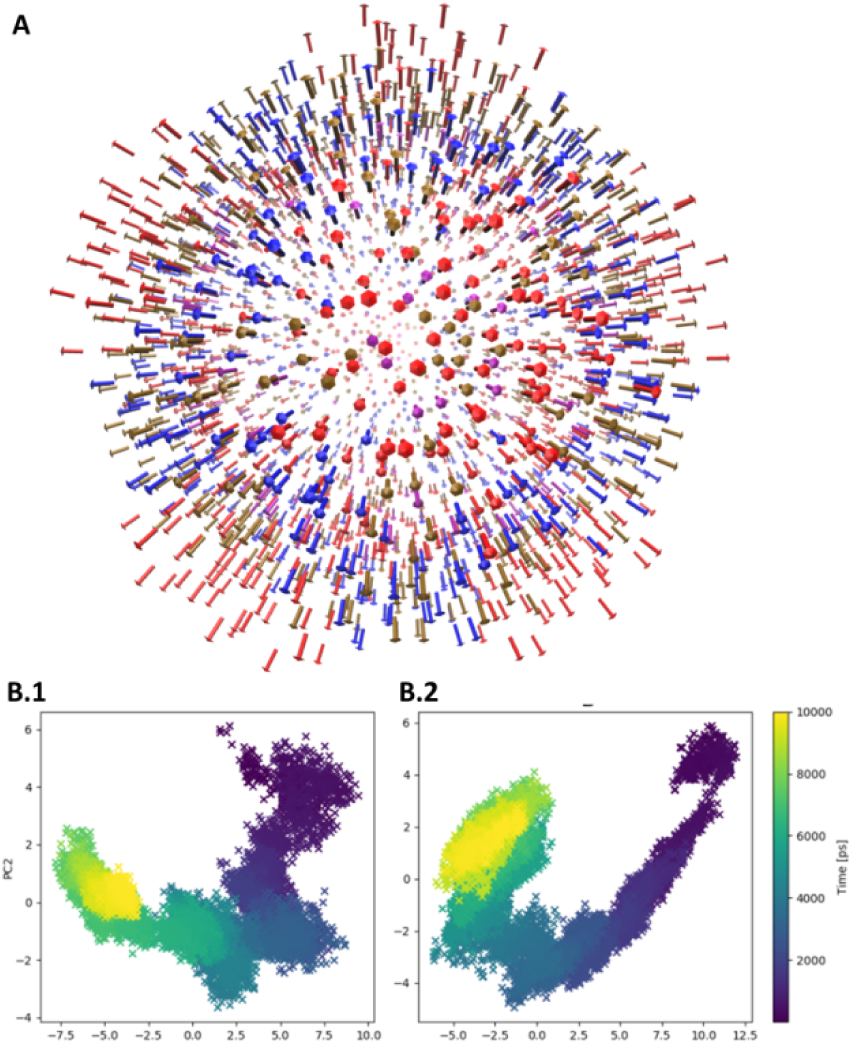
MODE-TASK visual outputs. (A) Eigenvectors of the expan sion mode projected as arrows onto the coarse-grained capsid, coloured by subunit: VP1-red; VP2-blue; VP3-brown; VP4-purple. (B) Dimension reduction by PCA for WT (B.1) and mutant (B.2), coloured per MD time point.

### Essential dynamics

P40L variation is disruptive to the stability of the renin-angiotensinogen system (RAS, 781 residues) (Brown *et al.,* 2017). We explored the essential dynamics of the complex using standard PCA for the wild type versus the mutant RAS MD trajectories. Figure 1B depicts the shifts in the energy landscape of the protein upon mutation.

## 5 Conclusion

Here, we present a novel and comprehensive downloadable software suite, MODE-TASK, which integrates a set of tools to analyse protein dynamics obtained from MD simulations as well as coarse grained elastic network models.

## Funding

This work is supported by the National Institutes of Health Common Fund under grant number U41HG006941 to H3ABioNet, the National Research Foundation (NRF) South Africa (Grant Number 93690) and the Scientific and Technological Research Council of Turkey (Grant Number 116F229). The content of this publication is solely the responsibility of the authors and does not necessarily represent the official views of the funders.

## Conflict of Interest

none declared.

